# A common neural axis links image distinctiveness and memorability

**DOI:** 10.64898/2026.11.10.765672

**Authors:** Igor Utochkin, Woohyeuk Chang, Edward Vogel, Edward Awh

**Author notes:** Correspondence: Igor Utochkin. I.U. and W.C. contributed equally to this work.

## Abstract

Stimulus-specific memorability, the tendency for people to remember and forget the same stimuli, is a powerful predictor of memory performance that is independent of the study and retrieval factors that have dominated memory research. Critically, memorability can be defined in two ways: “Hit-ability” (H-ability) and “false alarm-ability” (FA-ability), the probability of correctly recognizing studied items or falsely recognizing unstudied items, respectively. Although past work has suggested that these are dissociable facets of memorability, challenging single-factor explanations, we present neural and computational evidence that they share a *common neural axis*. The neural signature of *high H-ability* matched the signature for *low FA-ability*, showing that these parameters have inverse relationships with a common factor. Moreover, we show that *image distinctiveness* predicts the flipped relationship between H-ability and FA-ability in both behavior and neural activity, providing a unifying explanation of stimulus-specific memorability.

## INTRODUCTION

For centuries, the dominant models of long-term memory have focused on study and retrieval conditions as the key drivers of memory performance. More recent work, however, has focused on *stimulus-specific memorability*, the finding that a compelling amount of variance in performance can be predicted based on the stimuli themselves, independent of study and retrieval conditions^1,2,3,4^. Importantly, memorability can be broken down into two aspects: Hit-ability (H-ability) refers to an item’s tendency to be correctly recognized when it is studied. False alarm-ability (FA-ability) refers to the probability that an item will be erroneously recognized when it was not studied. Remarkably, in behavioral tests of memory, these two aspects of stimulus memorability often show small or negligible correlations (^5,6,7,8^; but see^9^), posing a challenge for single-factor explanations. Nevertheless, here we show that H-ability and FA-ability have inverse relationships with a *common neural axis,* and that *image distinctiveness* may explain this shared axis.

We collected a dataset of 145 electroencephalogram (EEG) recording sessions from human participants as they memorized and recognized visual images of objects from a large set of 2,400 pictures^10^. We obtained H- and FA-ability scores for all of these images in a large online experiment with 989 participants. First, we demonstrate robust decoding of both H-ability and FA-ability based on the scalp topography EEG activity evoked during both the study and test phases of a recognition memory experiment. Second, we show that *a common neural axis* captures the neural signatures of H-ability and FA-ability, such that the neural pattern associated with *high H-ability* resembles the neural pattern associated with *low FA-ability*. Thus, H-ability and FA-ability have mirror-imaged neural signatures, pointing to a single-factor explanation of memorability.

What is this single factor? Although past work has shown that H-ability and FA-ability can be modeled as biases along a common *familiarity* axis^7^, this cannot explain the flipped neural patterns that we observed for these parameters. Thus, we considered the potential role of image *distinctiveness*, a factor that has been highlighted in past formal models of stimulus-specific memorability^11,12,13,14^. To this end, we used artificial neural networks (ANNs) that are known to track human similarity judgements^15,16^ to quantify the psychological distinctiveness of every item in our image database. This computational estimate of distinctiveness was found to reliably predict behavioral measures of memorability, although this link was primarily driven by an inverse relationship between FA-ability and distinctiveness. Interestingly, the link between distinctiveness and memorability was strengthened when distinctiveness was defined based on the later layers of ANNs that capture semantic and conceptual features^17,18^. Critically, neural classifiers that were trained to decode distinctiveness also enabled robust decoding of H-ability and FA-ability. The neural pattern for higher distinctiveness aligned with the neural patterns for high H-ability and low FA-ability. Thus, distinctiveness may provide a unifying explanation for these two aspects of stimulus-specific memorability.

## RESULTS

### Relationships between H-ability, FA-ability, and image distinctiveness

We collected H-ability and FA-ability scores of 2,400 images of everyday objects from Brady et al.^10^ in an online experiment including 989 participants performing an old/new recognition task (Fig. 1a). Each image was tested for hit rate (as target) and false-alarm rate (as foil) in approximately 120 participants. In line with a number of previous demonstrations^5,6,7,8^, we found only a small negative correlation between z-transformed hit and false-alarm rates, z(H) and z(FA) (Spearman’s ρ = -0.155, p < 0.001; Fig. 1b), suggesting a substantial degree of behavioral independence.

**Fig. 1.**
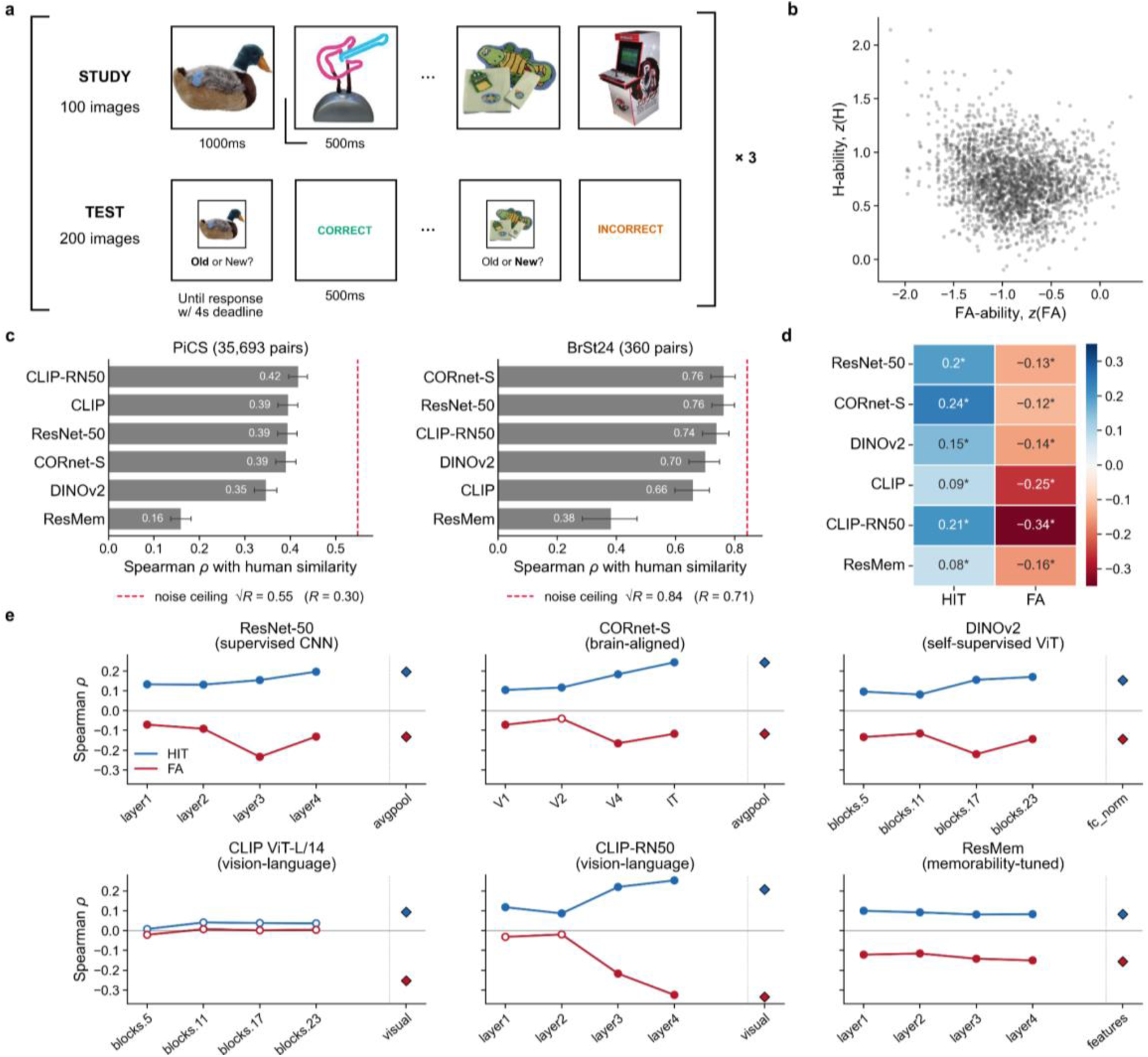
Relationship between H-ability, FA-ability and ANN-derived image distinctiveness. **a,** Old/new recognition task used to obtain item-level memorability scores for 2,400 object images from 989 online participants. Participants studied images and subsequently judged targets and foils as old or new. **b,** relationship between H-ability and FA-ability, across images. Each dot represents one image. **c,** Correlations between model-derived pairwise image similarities and human similarity ratings from two independent datasets (35,693 pairwise similarities across 1,200 objects from the PiCS database,^19^; 360 unique pairs from^20^) across six ANNs. Bars indicate Spearman’s ρ; error bars indicate 95% bootstrap confidence intervals; dashed lines indicate reliability-based noise ceilings. **d,** Spearman correlation between final embedding image distinctiveness and H-ability (blue) or FA-ability (red) for each ANN. **e,** Corresponding correlations across processing stages of each model; diamonds indicate final embeddings. In **d,e,** significance was assessed using two-sided permutation tests with Benjamini-Hochberg FDR correction across model, representation and memorability comparisons. Asterisks (**d)** and filled markers (**e**) indicate significant correlations (q < 0.05).

Despite this weak relationship, H-ability and FA-ability may nevertheless be linked with a common stimulus property. One candidate is *image distinctiveness*, which has been proposed to promote correct recognition and reduce false recognition by limiting confusability with other representations^21^. To quantify this property, we computed a measure of image distinctiveness using six ANNs spanning different architectures and training objectives (see Methods). We first confirmed that all six models captured human-like similarity structure (Fig. 1c): model-derived pairwise similarities correlated with human similarity ratings in two independent datasets (^20^; PiCS,^19^). For each model layer, we defined distinctiveness as 1 minus an image’s mean cosine similarity to all other images in the stimulus set. Distinctiveness was associated with both components of memorability, but in opposite directions. At the final embeddings, distinctiveness correlated positively with z(H) (ρ = 0.083 to 0.243) and negatively with z(FA) (ρ = -0.117 to -0.335) in all six models (all q < .001; Fig. 1d). The same pattern held across most intermediate representations, surviving FDR correction in 26 of 30 for H-ability and 23 of 30 for FA-ability (Fig. 1e). These associations generally strengthened at later processing stages, although depth profiles varied across models. Thus, more distinctive images tended to be both better recognized as targets and less often falsely recognized as foils.

Although z(H), z(FA), and image distinctiveness shared some variance, as suggested by their correlations, we did not find evidence for substantial multicollinearity between these variables (variance inflation factors did not exceed 1.2). That allowed us to consider H-ability, FA-ability, and distinctiveness as relatively independent factors in the models of neural activity in the recognition memory tasks. We performed a median split of the stimuli along each dimension such that each item was labeled as having “low” or “high” value in H-ability, FA-ability, and distinctiveness. These labels were further used as ground-truth stimulus values in the EEG analyses.

### H-ability and FA-ability are decodable in study and test phases

Motivated by past work showing that evoked EEG activity tracks stimulus memorability^23^, we sought to examine whether neural decoding could help to elucidate the links between H-ability and FA-ability. A dataset of 145 EEG recording sessions with participants performing various recognition memory tasks with the same images was used to determine the neural signatures of memorability during both study and test phases of the task. We first asked whether H-ability and FA-ability could be decoded based on evoked EEG activity (Fig. 2a). Both dimensions showed sustained decodability during study and test, emerging at approximately 150-200 ms after stimulus onset and remaining significant through the end of the epoch (FDR-corrected, q<.05; Fig. 2b-e). It is worth highlighting here that the memorability parameter estimates were obtained from a distinct group of subjects (collected online) from the ones that participated in the EEG recording sessions. Thus, the orderly correspondence between the behavioral parameter estimates and the neural signatures of memorability shows that these are truly *item-specific* effects rather than effects driven by situational changes in memory strength or decision challenges.

**Fig. 2.**
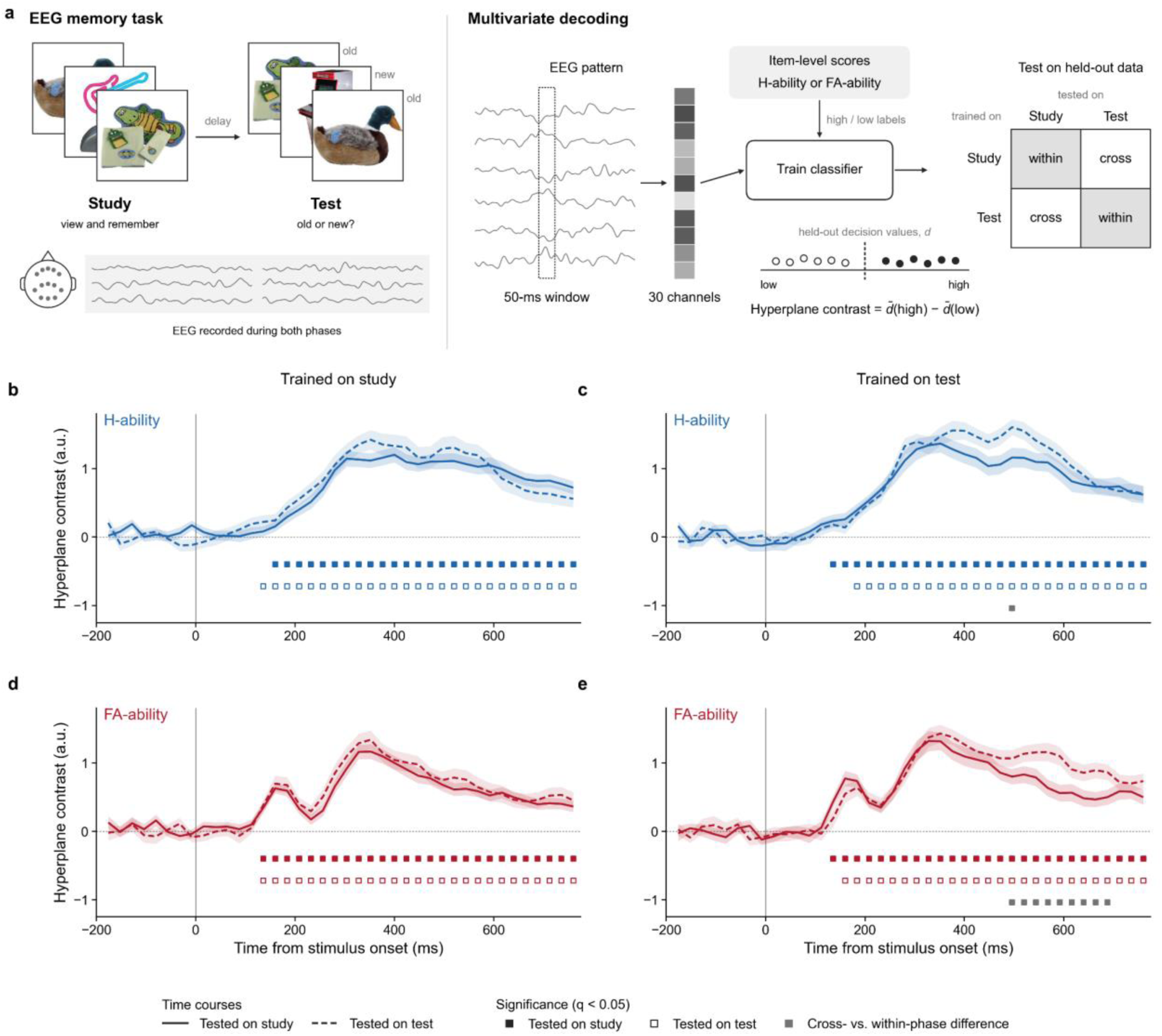
Neural signatures of H-ability and FA-ability emerge at encoding and generalize across memory phases. **a**, Schematic of the EEG recognition-memory task and multivariate decoding procedure. EEG was recorded during both study and test phases. Classifiers trained to distinguish high- and low-memorability images, based on independently obtained item-level H-ability or FA-ability estimates, were evaluated within and across memory phases. Decoding was quantified using the hyperplane contrast, defined as the difference in mean signed decision values between high- and low-memorability images. **b-e,** Hyperplane contrast over time for H-ability (**b,c**; blue) and FA-ability (**d,e**; red), with classifiers trained on study (**b,d**) or test (**c,e**) trials. Solid and dashed lines indicate classifiers tested on study and test trials, respectively. Filled and open squares indicate contrasts significantly different from zero (two-tailed one-sample *t*-tests); grey squares indicate significant differences between within- and cross-phase decoding (two-tailed paired *t*-tests). All tests were FDR-corrected across post-stimulus time points (q < 0.05). Shading indicates ±1 standard error of the mean (SEM) across sessions. Sample sizes were n = 125 sessions for within-phase study decoding, n = 80 for within-phase test decoding and n = 60 for cross-phase decoding.

Next, we found that these neural signatures generalized across study and test phases: classifiers trained in one phase produced significant hyperplane contrasts when tested in the other, with a similar onset and duration. For H-ability, cross-phase decodability showed no reliable reduction relative to within-phase decodability in either direction (Fig. 2b,c). FA-ability also showed robust study-to-test generalization (Fig. 2d), although test-to-study generalization was reduced relative to within-test decodability during a later half of the epoch (500 - 700 ms from stimulus onset; Fig. 2e). One possible explanation of reduced generalization from test to study is that there may have been decision-related activity during test that was correlated with memorability (e.g., decision confidence likely varied with memorability) but absent during study. Indeed, the robust decoding observed during the study period rules out decision-related activity as a driver of the common neural signature of memorability across the study and test phases.

### The neural signatures of H-ability and FA-ability are independent from mnemonic status

Because H-ability and FA-ability describe an item’s tendency to be recognized when presented as a target or foil, respectively, we asked whether their neural signatures during memory test could be distinguished from the neural representation of the item’s actual mnemonic status (old or new). We therefore performed representational similarity analysis (RSA) jointly modeling H-ability, FA-ability, and mnemonic status (Fig. 3a).We found that, even though the mnemonic status was a powerful predictor of EEG signals at memory test, H-ability and FA-ability both explained reliable unique variance in the sustained temporal window starting at approximately 200 ms (Fig. 3b). Thus, even during test phases, the neural effects associated with H-ability and FA-ability could not be reduced to whether the current item was objectively old or new.

**Fig. 3.**
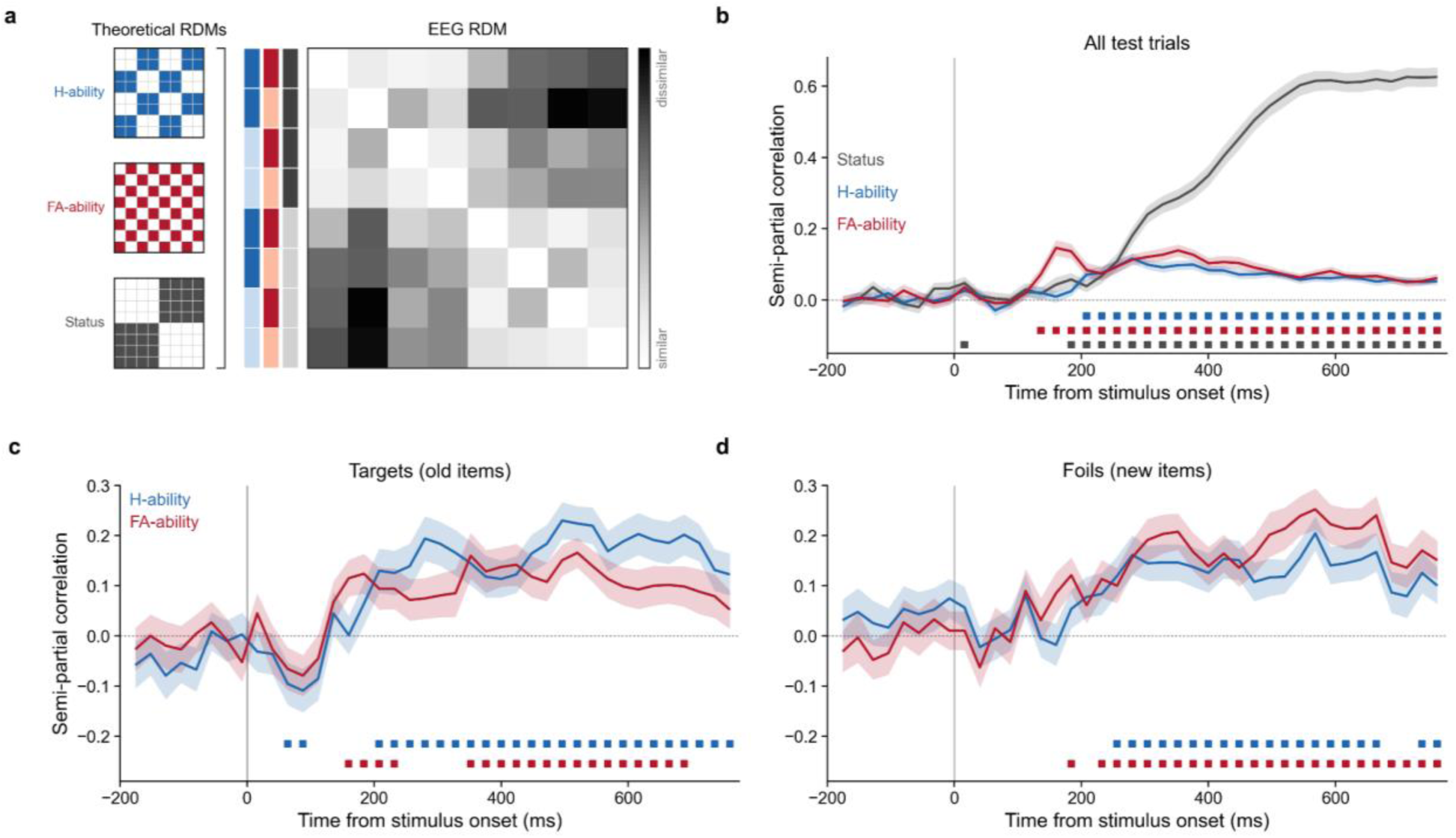
Neural signatures of H-ability and FA-ability are independent of mnemonic status. **a**, Theoretical representational dissimilarity matrices (RDM) for H-ability, FA-ability and mnemonic status (old/new) and empirical EEG RDM, defined across the eight conditions of the 2 x 2 x 2 factorial design. Empirical RDMs were computed using cross-validated Mahalanobis distances. Theoretical RDMs were entered simultaneously to obtain semipartial rank correlations indexing the unique contributions of each factor. **b,** Test-phase RSA showing the unique contributions of H-ability (blue), FA-ability (red) and mnemonic status (grey). **c,d**, RSA restricted to target (**c**) or foil (**d**) trials, with H-ability and FA-ability entered simultaneously. Conditions were defined by H-ability x FA-ability. Squares indicate semipartial correlations significantly different from zero (two-tailed one-sample *t*-tests, FDR corrected across post-stimulus time points, q < 0.05). Shading indicates ±1 SEM across sessions. Sample size was n = 80 sessions for all analyses.

We next asked whether these effects were specific to the mnemonic role with which each measure is defined. For example, H-ability might contribute primarily on target trials, whereas FA-ability might contribute primarily on foil trials. To test this, we ran RSA for the target and foil trials separately and included only H-ability and FA-ability theoretical RDMs into the linear model. We found that both dimensions keep their unique contributions into both target and foil trials (Fig. 3c,d). Thus, the neural signatures of H-ability and FA-ability are not driven by mnemonic status.

### A shared neural axis for H-ability and FA-ability

Although H-ability and FA-ability each explained unique variance in EEG activity beyond mnemonic status, this does not show that they rely on distinct neural representations. We therefore asked whether there is a shared neural axis between H-ability and FA-ability by examining whether the neural signature of one parameter generalized to the other. For example, we examined whether a model trained to discriminate between high and low H-ability could discriminate between stimuli sorted on the basis of FA-ability (and vice versa). Indeed, we found evidence that H-ability and FA-ability are linked by a *common neural axis.* We observed robust generalization of the neural patterns for H-ability and FA-ability during both study and test phases, but with an *inverted* mapping between the two dimensions. Thus, the neural signature of *high* H-ability resembled the neural signature of *low* FA-ability, and the neural signature of *low* H-ability resembled the neural signature of *high* FA-ability (Fig. 4a-d). The within and cross-dimensional contrasts diverged significantly from approximately 150 ms after stimulus onset in all four analyses, with the divergence sustained across the remainder of the post-stimulus interval. Thus, H-ability and FA-ability are reliably tracked by a common neural axis, but with opposite polarity. This robust generalization of neural patterns for H-ability and FA-ability makes a striking contrast with behavioral data that reveal only weak correlations of H-ability and FA-ability at the item level.

**Fig. 4.**
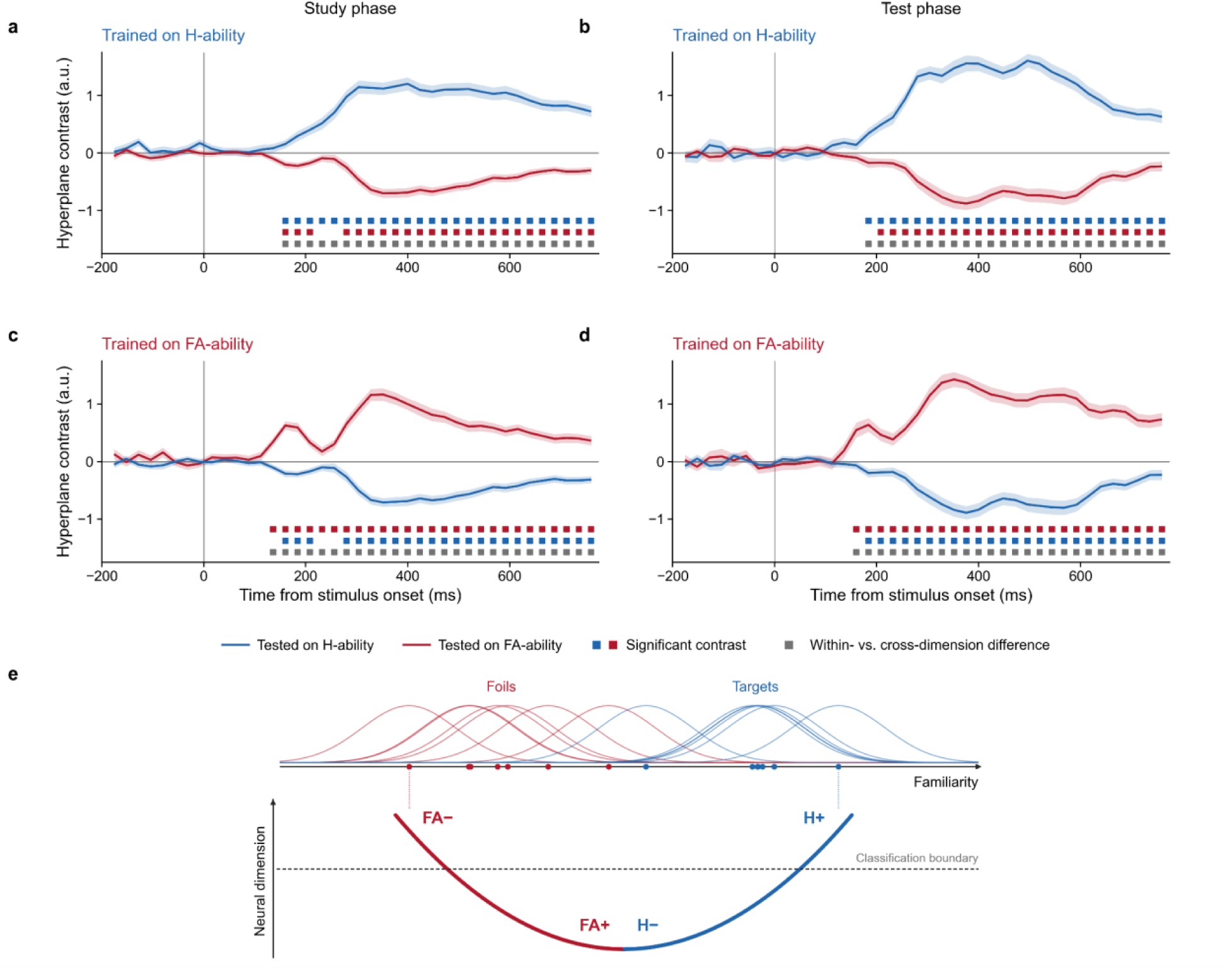
H-ability and FA-ability share a neural axis with opposite polarity. **a**–**d**, Cross-dimensional decoding of H-ability and FA-ability during study and test. Classifiers trained to distinguish high from low H-ability were tested on H-ability (blue, within-dimension) and FA-ability (red, cross-dimension) during study (**a**) and test (**b**). Classifiers trained on FA-ability were tested on FA-ability (red, within-dimension) and H-ability (blue, cross-dimension) during study (**c**) and test (**d**). Negative hyperplane contrasts indicate a reversal of the high- and low-memorability categories relative to the trained classification axis. Blue and red squares indicate contrasts significantly different from zero (two-tailed one-sample *t*-tests); grey squares indicate significant differences between within- and cross-dimension decoding (two-tailed paired *t*-tests). All tests were FDR-corrected across post-stimulus time points (q < 0.05). Shading indicates ±1 SEM across sessions. Sample sizes were n = 125 sessions for study and n = 80 for test. **e,** Hypothetical nonlinear relationship between familiarity and a shared neural dimension. Illustrative familiarity distributions for foils (red) and targets (blue) are shown along a common familiarity axis. A U-shaped mapping produces similar neural responses between low FA-ability (FA-) and high H-ability (H+) and between high FA-ability (FA+) and low H-ability (H-). The dashed classification boundary illustrates how such a mapping could produce the opposite polarity in cross decoding results in **a-d**.

What is the nature of the common neural axis that is shared by H-ability and FA-ability? Critically, even though past work has shown that H-ability and FA-ability can be precisely modeled as biases along a common *familiarity* axis^7^, the mirror-imaged neural pattern that we observed for these parameters is the opposite of what would be predicted by a familiarity account that predicts parallel increases in H-ability and FA-ability as familiarity increases (Fig. 4e). Thus, familiarity cannot explain the flipped neural pattern for H-ability and FA-ability. We also considered whether these neural patterns could be explained by confidence; after all, higher H-ability and lower FA-ability should both lead to higher confidence during testing. But we can also rule out confidence, because the same neural patterns are observed during *study* when observers are not even making any decisions.

### Distinctiveness as the common neural axis linking H-ability and FA-ability

What is the best explanation of the common neural axis shared by H-ability and FA-ability? Our behavioral analyses identified image distinctiveness as one candidate, with more distinctive images showing higher H-ability and lower FA-ability (Fig. 1d,e). We therefore examined whether the neural signature of ANN-derived image distinctiveness aligned with the neural signatures of these two memorability dimensions.

First, we found that distinctiveness could be reliably decoded from EEG activity during both study and test phases, as illustrated by the CLIP-RN50 time courses (Fig. 5a). This finding generalized across all six ANN-derived distinctiveness measures, which showed positive mean decoding contrasts during both phases (Fig. 5b). We next tested whether the neural signature of distinctiveness generalized to H-ability and FA-ability using cross-training analyses. Classifiers trained on CLIP-RN50 distinctiveness showed positive generalization to H-ability and negative generalization to FA-ability during both study and test, consistent with the opposing neural signatures observed in Figure 4 (Fig. 5c). The mean contrasts across all six ANNs showed a similar overall pattern, although the magnitude of generalization varied across models and was generally weaker for H-ability than for FA-ability, except for CLIP ViT-L/14, whose distinctiveness generalized negatively to H-ability (Fig. 5d). Together, these findings suggest that image distinctiveness may explain the common neural axis underlying H-ability and FA-ability.

**Fig. 5.**
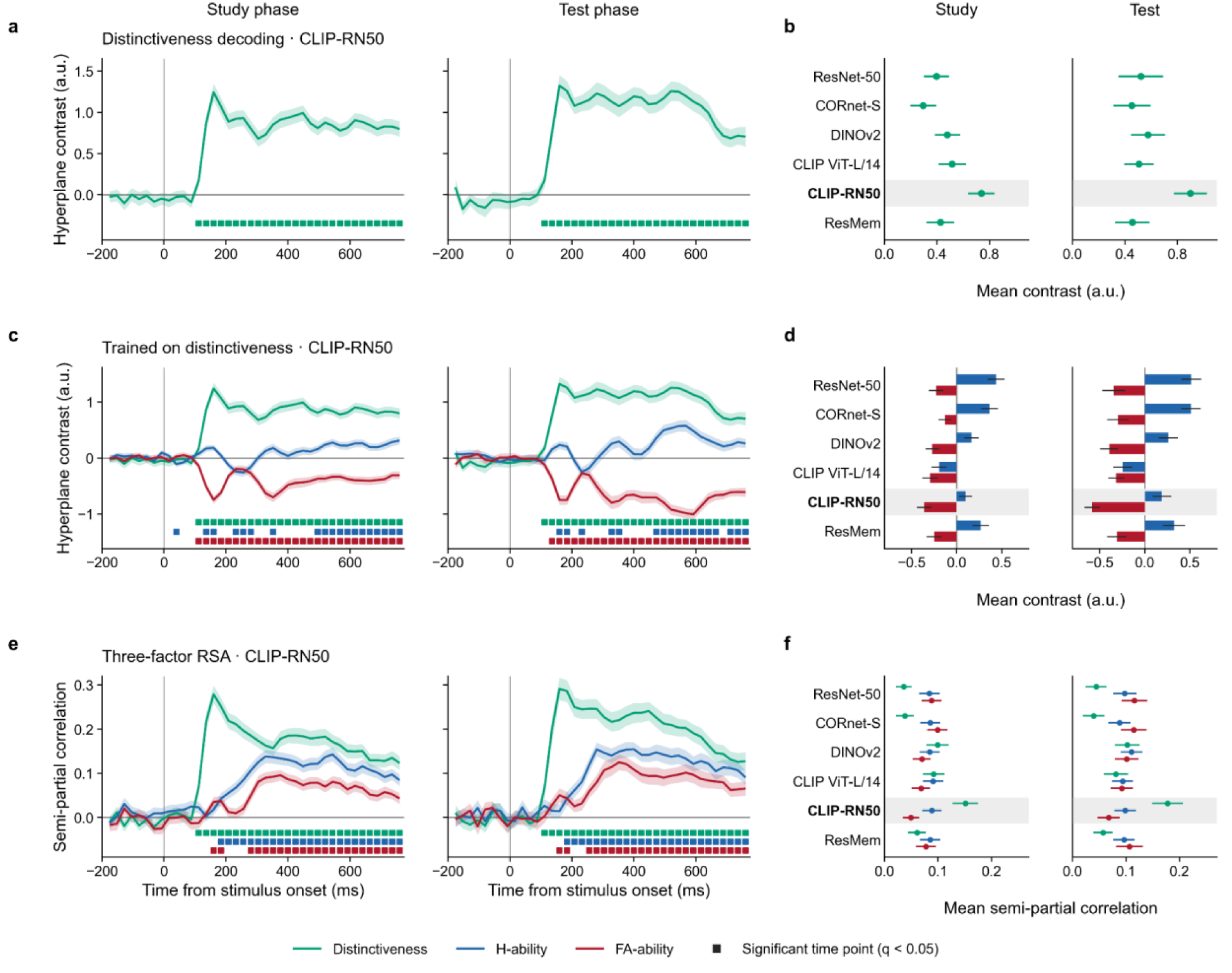
Distinctiveness shares a neural axis with H-ability and FA-ability but does not fully account for them. **a,c,e,** Time courses for CLIP-RN50 during study and test. **a**, Hyperplane contrast for high versus low image distinctiveness. **b**, Mean distinctiveness decoding across six ANN-derived measures. **c**, Cross-dimensional decoding with classifiers trained on distinctiveness and tested on distinctiveness (green), H-ability (blue) or FA-ability (red). **d**, Mean cross-dimensional decoding to H-ability and FA-ability across six ANNs. **e**, Three-factor RSA showing the unique contributions (semipartial correlations) of H-ability (blue), FA-ability (red) and distinctiveness (green), with all three theoretical RDMs entered simultaneously. **f**, Mean semipartial correlations across six ANNs. In **b,d,f**, CLIP-RN50 is highlighted, and values were averaged within sessions over the post-stimulus interval (0 - 800ms) before calculating group means. Squares in **a,c** indicate hyperplane contrasts significantly different from zero (two-tailed one-sample *t*-tests); squares in **e** indicate semipartial correlations significantly greater than zero (one-tailed Wilcoxon signed-rank tests). All tests were FDR-corrected across post-stimulus time points (q < 0.05). Shading indicates ±1 SEM across sessions. Sample sizes were n = 125 sessions for study and n = 80 for test.

To extend these findings, we used RSA to examine whether H-ability, FA-ability, and distinctiveness explain unique variance in EEG activity. We constructed empirical RDMs defined by the factorial combination of H-ability, FA-ability, and distinctiveness and examined all three theoretical models within a single RSA analysis. H-ability, FA-ability, and distinctiveness each showed significant positive semipartial correlations with the empirical RDM during both study and test phases (Fig. 5e,f). H-ability and FA-ability showed significant unique contributions beginning at approximately 150 ms after stimulus onset and persisting through the end. In contrast, the magnitude and temporal profile of the unique distinctiveness contribution varied across ANN models, with CLIP-RN50 showing the largest effect and a pronounced peak at approximately 200 ms. Together, these results indicate that distinctiveness captures a reliable portion of the neural variance, but H-ability and FA-ability also continue to explain unique variance. Thus, distinctiveness provides an attractive explanation for the common neural axis that tracks both facets of stimulus-specific memorability. That said, further work will be needed to characterize the unique remaining neural variance that is accounted for by H-ability and FA-ability.

## Discussion

People have a strong tendency to remember and forget the same images, independent of the study and retrieval conditions that have dominated memory research, a phenomenon that has been labeled stimulus-specific memorability. Critically, past work examining H-ability and FA-ability has shown that these parameters are largely uncorrelated at the item level, posing a challenge for single-factor explanations. Nevertheless, we present neural evidence that a single factor, image distinctiveness, may provide a unifying explanation of both these aspects of memorability. First, cross-training analyses showed that a neural model of H-ability generalized to stimuli sorted based on FA-ability, such that high H-ability resembled low FA-ability and low H-ability resembled high FA-ability. Thus, a single neural axis captures both aspects of memorability. Further work suggested that distinctiveness may explain this shared neural axis.

We acquired estimates of image distinctiveness with a diverse set of artificial neural networks (ANNs), and found that higher distinctiveness predicted lower FA-ability and somewhat higher H-ability (although the latter effect was modest, in line with past work:^12,7^). Moreover, image distinctiveness was decodable from stimulus-evoked EEG activity, and the resulting neural model generalized to H-ability and FA-ability. Items with high distinctiveness produced the same neural pattern as those with high H-ability or low FA-ability. Thus, distinctiveness provides a clear explanation for both behavioral measures of memorability and the flipped neural signatures of H-ability and FA-ability.

A virtue of our dataset is that it included stimulus-evoked activity from both the study and the test phases of a recognition memory task (whereas prior neural work did not separate study and test phases or did not require memorization of the stimuli at all;^22,23,24,25^).

Conveniently, we observed robust decoding that generalized across both study and test phases, even though the study phase did not include any testing or retrieval activity. Thus, the observed neural signature of memorability is not explained by mnemonic status (studied or not studied) or decision processes (e.g., confidence) that could be confounded with stimulus memorability.

Another novel contribution of our study is that it is the first to directly compare the neural signatures of H-ability and FA-ability rather than combining them into a single memorability metric (e.g., hit rate minus false alarm rate). Separating H-ability and FA-ability thus allowed us to gain further insight into how these parameters may be related. In this context, it is worth noting the apparent contradiction between the finding that H-ability and FA-ability are only weakly negatively correlated at the item level, even though we find relatively clear evidence that H-ability is the “opposite” of FA-ability, both in terms of neural signature and the relationship with image distinctiveness. While this is a counterintuitive constellation of findings, note that the flipped neural pattern observed for H-ability and FA-ability is at the *aggregate* level, while the almost negligible negative correlation between H-ability and FA-ability was at the *item* level.

Even though we observed clear evidence that the neural signatures of H-ability and FA-ability are flipped, this does not entail a strong negative correlation at the within-item level. In other words, the shared neural axis implies that similar neural states emerge when observers encounter highly H-able or low FA-able items, but this does not mean that these states are caused by the same stimuli. Finally, while we present convincing converging evidence that distinctiveness may explain the flipped relationship between H-ability and FA-ability, the fact remains that H-ability and FA-ability both explained unique variance in our RSA analysis of those factors and distinctiveness. Thus, more work is needed for a comprehensive understanding of the factors that drive H-ability and FA-ability.

Our working hypothesis is that distinctiveness may explain the shared neural axis for H-ability and FA-ability. Importantly, we ruled out familiarity as a unified explanation, because that predicts an alignment between high H-ability and high FA-ability (the opposite of what we observed). Nevertheless, we are not minimizing the importance of familiarity for models of memorability. Familiarity (or general memory “strength”) has a long history of success in formal models of memory^26,27,28,29,30^. For instance, Utochkin et al^7^ precisely predicted memory performance in a 2-alternative forced choice test as a function of the H-ability of the target and the FA-ability of the foil. Thus, while familiarity may not explain the shared neural axis we have observed with EEG, it is an important part of a broader understanding of stimulus-specific memorability. To illustrate this point, Figure 4e depicts our hypothesis regarding the relationships between H-ability, FA-ability, distinctiveness, and familiarity. The horizontal axis represents familiarity. H-ability and FA-ability are viewed as positive and negative biases on the familiarity axis, respectively. The vertical axis represents a shared neural dimension, which we hypothesize as distinctiveness. Thus, images with high H-ability will be positioned towards the right extreme of the familiarity axis, while items with low FA-ability will be biased towards the left edge of the familiarity axis. We propose a non-monotonic relationship between distinctiveness and familiarity, such that higher distinctiveness is predicted for *both* items with high H-ability and items with low FA-ability. By contrast, items with low H-ability or high FA-ability should fall closer to the center of the familiarity axis. Thus, this illustration captures how distinctiveness can explain the flipped neural signatures of H-ability and FA-ability, while familiarity provides clear predictions for which items will be recognized as “old” and “new”.

Our proposal that distinctiveness provides a unifying explanation for H-ability and FA-ability falls in line with several influential computational models of memory^31,21^ and neurophysiological models of mnemonic pattern separation in the brain^32^. Moreover, some work has highlighted distinctiveness as an explanation of memorability per se, arguing that contextual distinctiveness allows easier retrieval of studied items and easier rejection of lures^12,14^. Although we obtained a “global” measure of distinctiveness by measuring each item’s average similarity to all other items in a large image database, even the small subsets that were presented to each subject were likely to replicate the statistical structure (and thus distinctiveness landscape) of the broader image database^14^. This correspondence between distinctiveness scores in small subsets compared to the overall list of images may explain why our global measures of distinctiveness were robustly decodable based on the smaller sets of images that each observer viewed. As such, when we split our stimuli into high and low distinctiveness groups relative to the stimulus set, we were able to find evidence for both correlation with behavioral measures of H-ability and FA-ability and reliably decode distinctiveness based on EEG activity. Thus, our parsimonious explanation of stimulus-specific memorability resonates with extant proposals.

Centuries of memory research have focused primarily on study and retrieval conditions for predicting memory performance. By contrast, stimulus-specific memorability explains strong variation in memory performance based only on the memory items themselves. Although past work has argued that memorability may be explained by distinct H-ability and FA-ability parameters, our findings suggest that these two facets of memorability share a common neural axis that is based on image distinctiveness. Thus, a sharper awareness of an item’s position within the global context of memory may provide important leaps in our ability to understand and predict memory performance.

## Methods

### Online memorability scoring

#### Participants

To obtain memorability scores online, we recruited participants via Prolific (www.prolific.com;^48^) and compensated at a rate of $10.00 per hour. Using Prolific’s built-in filters, eligibility was restricted to residents of the United States, aged 18 to 35, with normal or corrected-to-normal vision, no ongoing mental health condition or illness, and an approval rate above 99%. A total of 1088 participants completed the study. Ninety-nine participants were excluded for achieving below 55% accuracy in at least one block, yielding a final sample size of 989 participants. All participants provided informed consent before participation. The study was granted approval by the institutional review board at the University of Chicago.

#### Stimuli and procedure

Data collection was run on Pavlovia.org, a cloud-based platform for hosting psychophysical experiments. We used 2,400 images of category-unique real-world objects from the “Massive Memory” stimulus set (Brady et al.^10^). We created 48 lists consisting of 600 quasi-randomly sampled images (300 images were used as memory targets and 300 as foils in each list). Our quasi-random sampling algorithm guaranteed that all images appeared an equal number of times as target and foil items across lists. Each list was assigned to at least 20 participants. For each participant, we randomly shuffled the item order and split it into three subsets of 100 targets and 100 foils used for three separate blocks of the old/new memory task. In the study phase of each block, 100 target images were shown sequentially, each for one second with an interstimulus interval of 0.5 seconds. In the test phase, these 100 targets were randomly intermixed with 100 foils and presented one at a time. Participants had to respond to each test item whether it was old or new. At the beginning of the experiment, participants received a short practice block including 10 targets and 10 foils. For each memory list, data collection continued until at least 20 participants achieved above 55% accuracy. The hit and false alarm rates, p(H) and p(FA), for each test item were defined as the proportion of participants who responded “old” to that item, respectively.

#### H-ability and FA-ability estimates

For each image, we measured the hit rate and false-alarm rate as the proportion of participants who answered “old” to this image when this image was presented as a target or foil, respectively. The hit and false-alarm rates were then *z*-transformed to obtain *z*(H) and *z*(FA) indexes that can be conceptualized as the mean locations of item-specific distributions on a signal-detection familiarity axis when this item is a target or a foil^7^. In other words, *z*(H) and *z*(FA) show how much familiarity on average an item in a given mnemonic status causes across participants, which determines the H- and FA-ability of this item. For the purposes of subsequent EEG analysis, we split items into contrast categories of low or high H- and FA-ability via median split. The items with *z*(H) below the median were labeled as having “low H-ability”, whereas items with *z*(H) above the median as having “high H-ability”. The same thresholds applied to *z*(FA) yielded categories of low and high FA-ability.

### ANN-derived image distinctiveness

#### Model selection and feature extraction

To test whether the representational structure underlying memorability generalizes across different forms of visual computation, we extracted image features from six pretrained artificial neural networks (ANNs) that differed in architecture and training objectives. These included two supervised convolutional neural networks trained on object categorization (ResNet-50,^33^; CORnet-S,^34^), a self-supervised vision transformer trained without category labels (DINOv2,^35^), two vision-language models trained with contrastive image-text objectives (CLIP ViT-L/14 and CLIP-RN50,^36^), and a network trained directly to predict human memorability scores (ResMem,^37^). This model set allowed us to test whether findings generalized across architectures and training objectives. For each model, we extracted features from representative stages along its processing hierarchy. The specific layers and representations used are summarized in Extended Data Table 1.

#### Validation against human similarity judgments

To assess whether the representational spaces of the six models captured structure present in human similarity judgments, we compared model-derived similarities with two independent behavioral datasets: pairwise similarity ratings of real-world object images from Brady and Störmer^20^ (360 pairs) and validation ratings from the PiCS database (^19^; 35,693 pairs across 1,200 images from 20 categories). PiCS ratings were reverse coded from dissimilarity so that higher values indicated greater similarity in both datasets. For each image pair, we computed cosine similarity between model embeddings and correlated these values with mean human ratings using Spearman’s ρ. Because measurement noise limits the maximum observable model– human correlation, we estimated a reliability-based ceiling using the published reliability for Brady and Störmer and reliability estimated from trial-level PiCS ratings while accounting for unequal numbers of ratings across image pairs. Confidence intervals were obtained from 5,000 bootstrap resamples, resampling images for PiCS and image pairs for Brady and Störmer.

#### Image distinctiveness

For each selected layer of the six ANN models, we extracted a feature embedding for every image in the stimulus set using the THINGSvision toolbox^38^. Within each model-layer representation space, we computed pairwise cosine similarity between every image and all other images in the set. For each image, mean similarity was defined as its average cosine similarity to all other images, excluding self-similarity. Image distinctiveness was then defined as 1 - mean similarity, such that images with lower mean similarity to the rest of the stimulus set had higher distinctiveness. Distinctiveness was computed separately for each selected layer of each model.

To quantify the relationship between ANN-derived distinctiveness and memorability, we computed Spearman correlations between image distinctiveness and *z*(H) or *z*(FA) across the 2400 images, separately for each selected representation stage of each ANN, including the final embedding. Significance was evaluated using 5,000 two-sided permutations, with Benjamini-Hochberg FDR correction (q <.05;^39^) across all model x representation x memorability comparisons.

### EEG experiments

#### Participants and datasets

Participants for EEG experiments were recruited from the University of Chicago and nearby communities. In total, 145 EEG recording sessions were collected from 104 unique participants. As our joint EEG dataset comprises eight separate experiments, some of the participants took part in more than one experiment. Overall, 81 participants were tested in one experiment, 13 in two, 6 in three, 1 in four, 2 in five, and 1 in six experiments. A median time interval between any two consecutive sessions with a returning participant was 179 days. Before the beginning of the study, all participants gave a written informed consent whose form was approved by the Ethics Committee of the University of Chicago. We excluded participant’s data from analysis if their overall accuracy at memory test was below 55% correct or if EEG artifact rejection rate due to eye movements, blinks, or recording noise was over 30%. We also excluded the data of participants who did not complete the experiment.

#### Stimuli and experimental procedures

The EEG data sets for our analysis were collected from a set of full-sample and pilot (small-sample) experiments originally designed to test various research questions about recognition memory. As such, there were multiple variations across study and test conditions including variations in memory list length (from 8 to 100 items), the speed of presentation at the study phase (1 item each 1,750 ms or 1 item each 250 ms), study-test delay, and the form of memory test (old/new task or 2-alternative forced choice). Extended Data Table 2 presents details of each experiment and explains which subsets of the data were eligible for our current analysis.

The participants of the EEG experiments were tested in a darkened room with their head resting on a chin rest. Stimuli were presented on a Gamma corrected LCD monitor with a spatial resolution of 1,920 x 1,080 pixels and refresh rate of 120 Hz. The monitor was set at a 75-cm distance from the participant. Participants used a standard computer keyboard to respond.

Stimulus presentation and response registration were run in PsychoPy v. 3.2.4 for Windows^40^. The general design of each experiment included multiple blocks consisting of a study and test phase spaced by a 6-sec delay. Targets and foils for each participant and each block were randomly sampled from the “Massive memory” stimulus set^10^. The images never repeated between blocks within one session. In the study phase, participants were sequentially shown target images (approx. 6.3° of visual angle in each dimension) at a screen center. The presentation of each item was preceded by a fixation cross (400-600 ms before the image onset) on a uniform gray background. Each item appeared for 800 ms during which the fixation cross stayed on the screen. Participants were encouraged to look at the fixation cross and not to blink or move their eyes until both the cross and the target disappear. During a 1,000-ms intertrial interval (ITI) following the target offset and preceding a next fixation cross onset, a blank white box matching the target image size was present. Participants could blink during the ITI.

In the test blocks of the old/new recognition task, tested targets were intermixed with the same number of foils and presented in a random order at the screen center. The first trial of the test phase started automatically six seconds after the last target from the study phase. The rest of the test trials were started by participants pressing a spacebar. At the beginning of each trial, participants saw a fixation cross for 400-600 ms followed by a test image. During the first 800 ms after the test image onset, the fixation cross stayed on the screen and participants could not respond. After that, the fixation cross changed to a question mark signaling the participants to respond whether the image was old or new by pressing an “z” or “/” button of a standard keyboard with their left or right hand, respectively. Button assignment to a response category was balanced across participants. The test trial ended after a response or 4,000 ms. Participants were encouraged to keep their gaze on the fixation cross or the question mark and not to blink throughout the trial. After the trial ended, a blank white square and an instruction “Press space for the next trial” were presented. The percentage of correct answers was presented after each practice or main block.

#### EEG and eye-tracking acquisition

We recorded EEG from 32 active Ag/AgCl electrodes (actiCHamp, Brain Products, Munich, Germany) mounted according to the International 10-20 system. They included Fp1/2, F3/4, F7/8, Fz, FC1/2, FC5/6, C3/4, Cz, CP1/2, CP5/6, P3/4, P7/8, Pz, PO3/4, PO7/8, O1/2, and Oz. Two electrodes were set on the right and left mastoids, and their average was used for offline re-referencing of EEG recorded from other sites. The ground electrode was set in the position Fpz. During preparation for recording, the EEG electrode impedance was set at or below 5 kOhms. Eye position was recorded using a desk mounted eye-tracking system EyeLink 1000 Plus (SR Research, Ottawa, Ontario, Canada). The eye-tracker was calibrated while participants followed five points on the screen with their gaze. Calibration accuracy was considered acceptable if the gaze position at the central point did not deviate from that point more than 0.5 degrees at the validation step. In addition to eye-tracking, electrooculograms (EOG) were recorded from five passive electrodes (horizontal EOG were recorded at the outer canthi of each eye, vertical EOG from above and below the right eye, ground was set below the left eye).

#### EEG preprocessing and artifact rejection

We performed EEG preprocessing and artifact rejection using EEGLAB^41^ with a standard pipeline from our laboratory (see detailed description in^42^). We epoched our EEG data from -200 to 800 ms relative to the stimulus onset both in the study and test phases. We rejected epochs where eye blinks or saccades with a length of over 0.5 degrees were detected according to eye-tracking. In cases when eye-tracking data was unavailable due technical reasons, rejection was made based on EOG data (potential saccades were flagged when a difference between HEOG or/and VEOG changed by over 20 μV within a 150-ms sliding window with a step of 10 ms).

### Multivariate EEG analyses

#### EEG decoding

We used a within-participant decoding framework^43^ using LogisticRegression in scikit-learn on Python^44^ and independently replicated with a logistic linear model in R. Analyses used EEG voltages from all electrodes except Fp1/2 from −200 to 800 ms relative to stimulus onset, baseline-corrected to −200 to 0 ms and averaged within 50-ms windows advanced in 25-ms steps. Trials were labeled as low or high in H-ability, FA-ability, and each ANN-derived distinctiveness measure using median splits of the corresponding item-level scores.

Because each image carried labels on multiple dimensions, within-phase cross-dimension decoding could otherwise use the same physical trials for both training and testing. We therefore split eligible within-phase trials into disjoint 70/30 training and test sets before trial binning, stratified by the joint combination of training and test labels. For cross-phase analyses, study and test trials were physically distinct, so all eligible trials from the respective phases were used. Within each pool, trials from the same class were averaged in bins of 20. At each time point, training and test data were standardized using the training-set mean and standard deviation. The procedure was repeated 1,000 times and decoding estimates were averaged across iterations.

Decodability was quantified using the hyperplane contrast^45,42,46^, defined as the difference between conditions in their mean signed distance from the fitted decision boundary. The magnitude of the contrast indexed decodability, whereas its sign indicated the alignment of the test representation with the trained decoding axis.

We performed all 16 train–test combinations among H-ability-study, H-ability-test, FA-ability-study, and FA-ability-test, capturing within- and cross-phase decoding as well as bidirectional H-ability–FA-ability cross-decoding. For each of the six ANN-derived distinctiveness measures, we additionally tested within-phase distinctiveness decoding and bidirectional cross-generalization between distinctiveness and H-ability or FA-ability. Cross-phase analyses were restricted to datasets containing both study and test phases; single-phase datasets contributed only to analyses of their available phase.

At each post-stimulus time point, hyperplane contrasts were tested against zero using two-tailed one-sample *t* tests across sessions. To assess generalization, generalized hyperplane contrasts were compared with the corresponding within-condition contrasts using two-tailed paired-samples *t* tests among sessions contributing to both analyses. Other planned contrasts between analyses were tested using the same paired-samples procedure. The Benjamini–Hochberg procedure controlled the false discovery rate at .05 across post-stimulus time points within each analysis or comparison.

#### Representational similarity analysis

We used representational similarity analysis (RSA), following Jones et al.^42^, to estimate the unique contributions of H-ability, FA-ability, and distinctiveness to EEG representational structure. RSA used the same electrodes and temporal averaging as the decoding analyses. For each participant, memory phase, time point, and ANN-derived distinctiveness measure, trials were assigned to eight conditions defined by the factorial combination of median-split H-ability, FA-ability, and distinctiveness (2 × 2 × 2). In the RSA testing the separability of between the neural signals of H- and FA-ability from mnemonic status, test-phase trials were split by the combinations of H-ability, FA-ability, and mnemonic status. This was complemented by separate RSAs within each mnemonic status, in which target and foil trials were assigned to four conditions defined by H-ability and FA-ability (2 x 2).

Empirical representational dissimilarity matrices (RDMs) were constructed using cross-validated Mahalanobis distance^46^ with Ledoit–Wolf covariance regularization^47^. Trials were randomly divided into two equal, condition-stratified partitions, and pairwise distances between the eight conditions were estimated across 1,000 random splits and averaged to obtain an 8 × 8 RDM at each time point.

Separate theoretical RDMs represented H-ability, FA-ability, and distinctiveness. Empirical and theoretical dissimilarities were rank-transformed, and the three theoretical models were entered simultaneously to obtain semipartial rank correlations indexing the unique variance associated with each factor after accounting for the other two. For the analyses of mnemonic status (Fig. 3), semipartial correlations were tested against zero at each time point from 0 to 800 ms using two-tailed one-sample *t*-tests across sessions. For the three-factor analyses with distinctiveness (Fig. 5), they were tested at each post-stimulus time point using one-tailed Wilcoxon signed-rank tests. In both cases, the Benjamimi-Hochberg FDR correction at .05 was applied across time points.

## Data availability

All data needed to reproduce the analyses and figures are openly available on Zenodo at https://doi.org/10.5281/zenodo.23751375. All analysis code is available at https://github.com/woohyeuk-leo-chang/mvMem.

## Notes

### Competing Interest Statement

The authors have declared no competing interest.

