## Extended Data Table for "A common neural axis links image distinctiveness and memorability"

**Extended Data and Supporting Tables**

### **Extended Data Fig. 1 | Item-level memorability distributions**

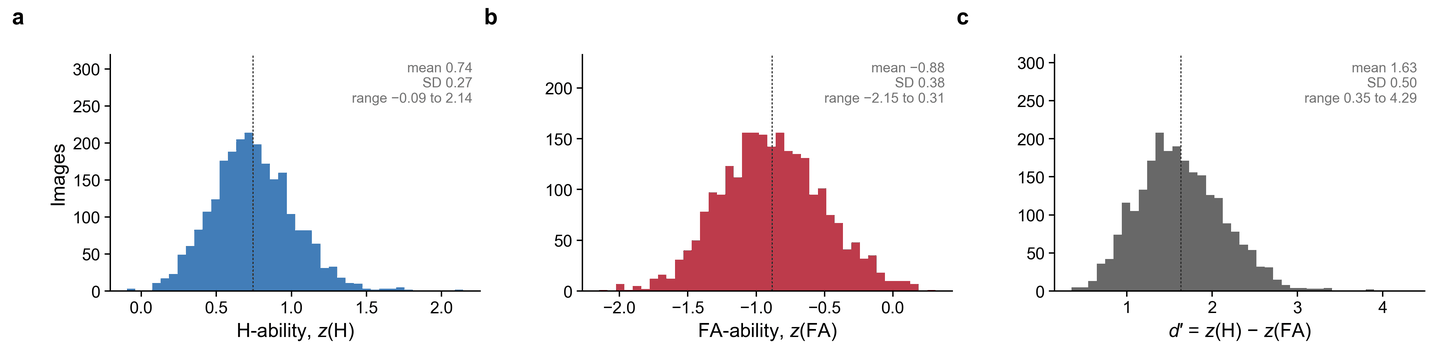

**Extended Data Fig. 1 | Item-level memorability distributions.** **a**–**c**, Distributions of H-ability, *z*(H) (**a**), FA-ability, *z*(FA) (**b**), and *d*′ = *z*(H) − *z*(FA) (**c**) across the 2,400 object images, from the online recognition task (*n* = 989 participants; each image was rated by a median of 123 participants as a target, range 118–133, and a median of 123 as a foil, range 117–134). Dashed lines, means; inset text, mean, s.d. and range.

### **Extended Data Fig. 2 | Distinctiveness decoding for all six ANN models**

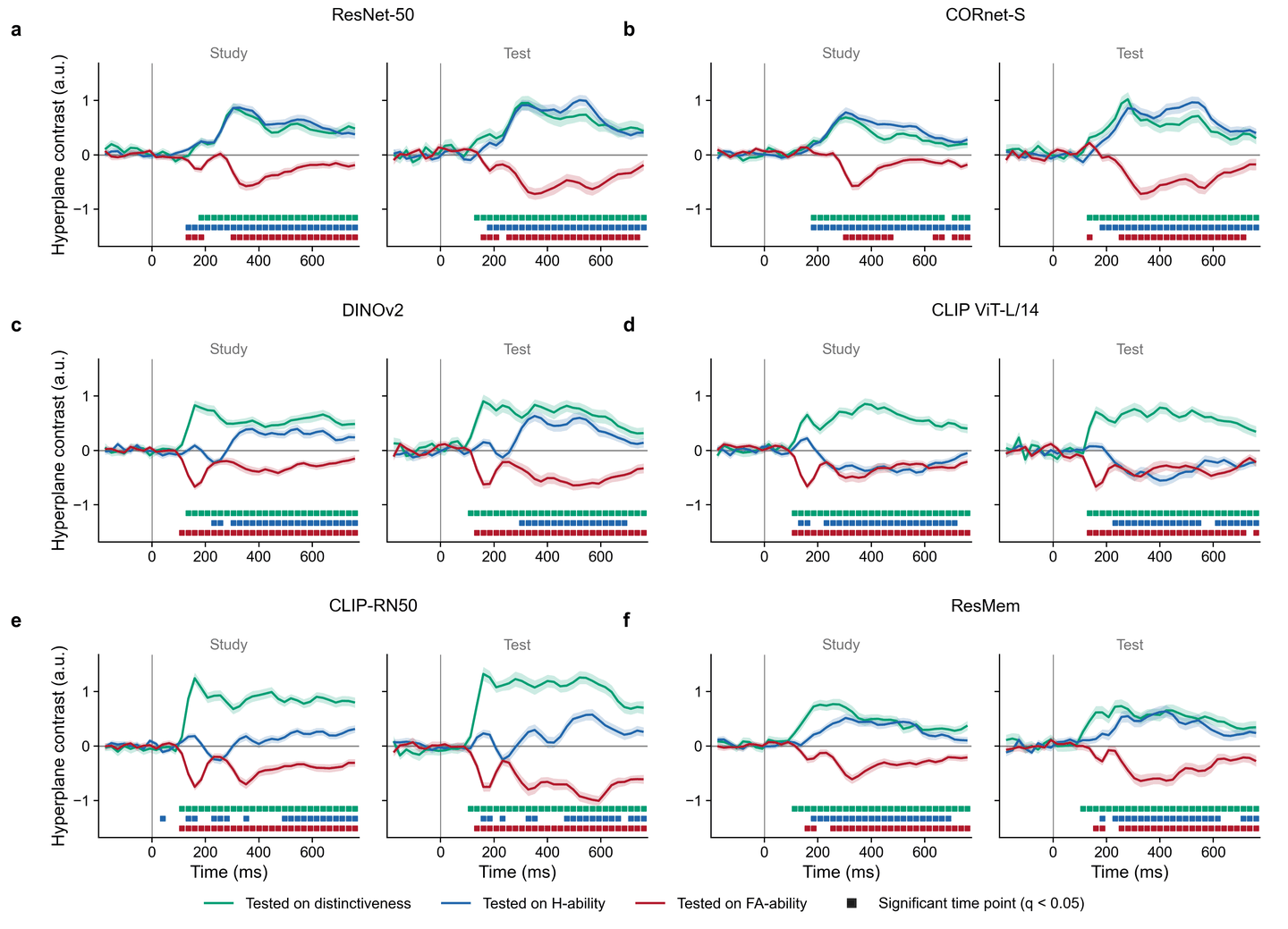

**Extended Data Fig. 2 | Distinctiveness decoding for all six ANN models.** **a**–**f**, For each ANN-derived distinctiveness measure, classifiers were trained to distinguish high- from low-distinctiveness images (median split) and tested on distinctiveness (green), H-ability (blue) and FA-ability (red) within the study (left) or test (right) phase. Lines, mean hyperplane contrast; shaded areas, ±1 s.e.m. across participants; squares, time points at which the contrast differed from zero (two-tailed one-sample *t*-tests, FDR-corrected across post-stimulus time points, *q* < 0.05). *n* = 125 participants (study) and 80 (test). CLIP-RN50 (**e**) is the example shown in Fig. 5a,c; six-model summaries are in Fig. 5b,d.

### **Extended Data Fig. 3 | Three-factor RSA for all six ANN models**

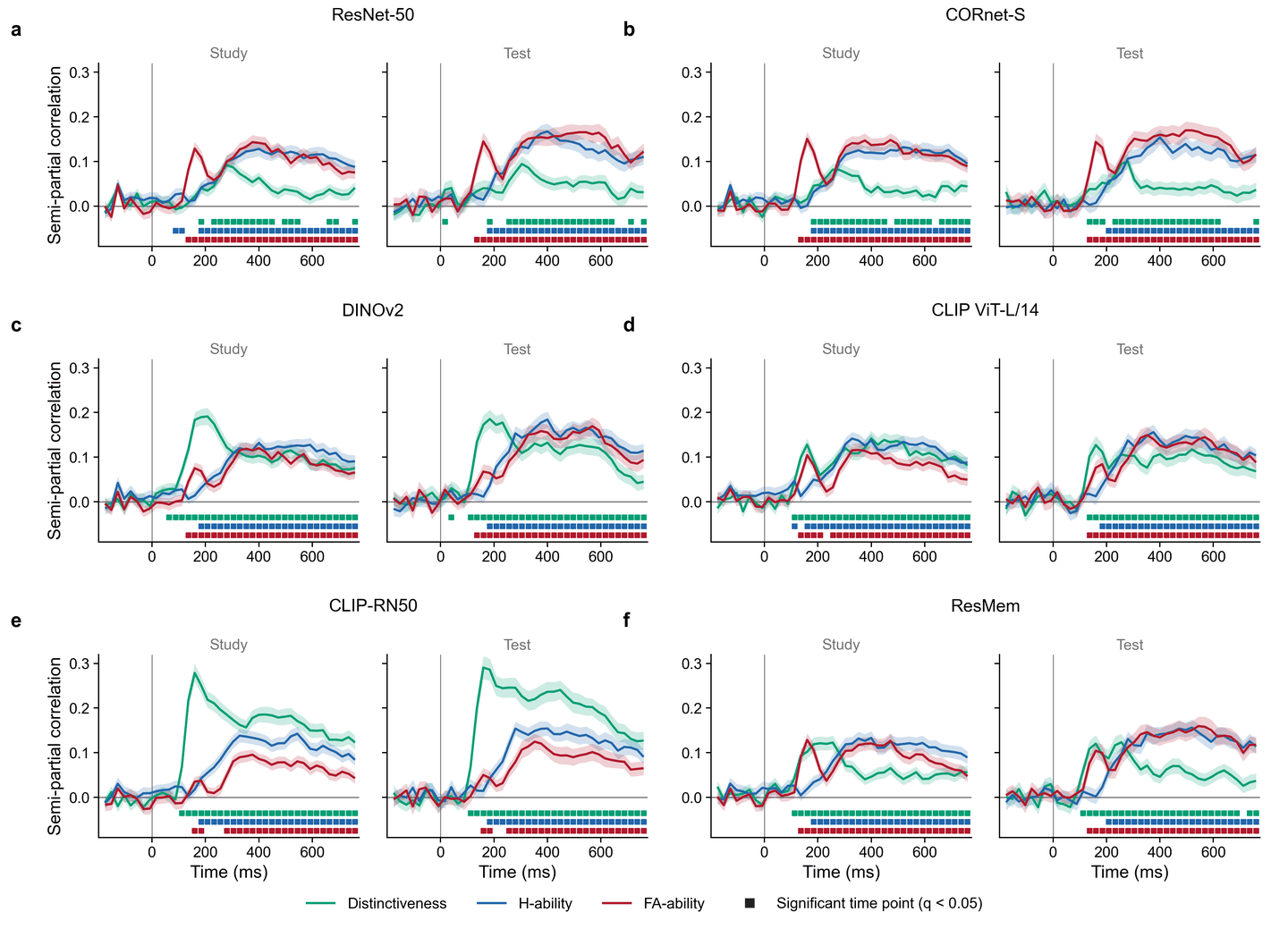

**Extended Data Fig. 3 | Three-factor RSA for all six ANN models.** **a**–**f**, Unique contributions (semi-partial rank correlations) of H-ability (blue), FA-ability (red) and distinctiveness (green) to the EEG representational geometry, from RSA with all three theoretical RDMs entered simultaneously (median-split 2 × 2 × 2 design), during the study (left) and test (right) phase, for each ANN-derived distinctiveness measure. Shaded areas, ±1 s.e.m. across participants; squares, semi-partial correlations exceeding zero (one-tailed Wilcoxon signed-rank tests, FDR-corrected across post-stimulus time points, *q* < 0.05). *n* = 125 participants (study) and 80 (test). CLIP-RN50 (**e**) is the example shown in Fig. 5e; six-model summaries are in Fig. 5f.

### **Supporting Table 2 | EEG dataset composition**

| **Experiment** | **Sessions (included)** | **Memory phases** | **Trials, study: total (per-session range)** | **Trials, test: total (per-session range)** | **List length(s)** | **Presentation rate (study): Stimulus duration + average ITI** | **Memory-test format** | **Brief experiment description, special notes** |
| --- | --- | --- | --- | --- | --- | --- | --- | --- |
| E1 | 20 | study + test | 6,546 (197–366) | 13,720 (617–751) | 8, 32, 64 | 800 + 1,500 | Old/New | The experiment tested list-length effects on ERP markers of old/new recognition |
| E2 | 20 | study + test | 18,788 (678–1101) | 16,564 (649–937) | 10, 32 | 800 + 1,500 | Old/New | The experiment tested list-length and primacy-recency effects on ERP markers of old/new recognition |
| E3 | 20 | study + test | 4,314 (191–236) | 4,437 (195–238) | 20 | 800 + 1,500 | Old/New | The experiment tested effects of interstimulus similarity of study and test lists on ERP markers of old/new recognition  Note: The experiment used images from two stimulus sets. The present analysis is limited to the trials where stimuli from Brady et al. (2008) were used, for which memorability scores are available |
| E4 | 4 | study | 1,962 (436–525) | — | 8, 32, 64 | 800 + 1,500 | 2-AFC | The experiment tested EEG signatures of recognition memory in 2-AFC |
| E5 | 30 | study | 17,355 (362–704) | — | 20 | 800 + 1,500 | 2-AFC | The experiment tested EEG signatures of recognition memory in 2-AFC |
| E6 | 6 | study | 3,458 (355–648) | — | 20 | 800 + 1,500 | 2-AFC | The experiment tested EEG signatures of recognition memory in 2-AFC |
| E7 | 25 | study | 25,220 (804–1169) | — | 20, 100 | 800 + 1,500 | 2-AFC | The experiment tested EEG signatures of recognition memory in 2-AFC |
| E8 | 20 | test | — | 11,246 (483–621) | 20 | 800 + 1,500,  250 + 0 | Old/New | The experiment tested effects of encoding speed on ERP markers of old/new recognition |
| **Total** | **145** |  | **77,643** | **45,967** |  |  |  |  |

### **Supporting Table 1 | ANN architectures and representations**

| **Model** | **Architecture and training objective** | **Weights / checkpoint** | **Representations (early → late; last = final embedding)** | **Output dimensions** | **Preprocessing** | **Pooling to one vector per image** |
| --- | --- | --- | --- | --- | --- | --- |
| ResNet-50 | Supervised CNN; ImageNet object classification (He et al., 2016) | torchvision resnet50, ImageNet weights IMAGENET1K_V2 (checkpoint resnet50-11ad3fa6.pth), pretrained=True, loaded via thingsvision | layer1, layer2, layer3, layer4, avgpool | 256, 512, 1,024, 2,048, 2,048 | thingsvision model-specific transforms | Conv stages: global average pooling; avgpool: as output |
| CORnet-S | Recurrent CNN aligned with the ventral visual stream; ImageNet classification (Kubilius et al., 2019) | thingsvision custom model cornet-s, pretrained=True | V1, V2, V4, IT, decoder.avgpool | 64, 128, 256, 512, 512 | thingsvision model-specific transforms | Areas: global average pooling; decoder.avgpool: as output |
| DINOv2 | Self-supervised vision transformer, ViT-L/14, LVD-142M (Oquab et al., 2023) | timm vit_large_patch14_dinov2.lvd142m, pretrained=True, via thingsvision | blocks.5, blocks.11, blocks.17, blocks.23, fc_norm | 1,024 (all five) | thingsvision model-specific transforms | Blocks: mean of patch tokens excluding CLS; fc_norm: pooled CLS token after final LayerNorm |
| CLIP ViT-L/14 | Vision–language model, contrastive image–text training (Radford et al., 2021) | thingsvision custom model clip, variant ViT-L/14 (OpenAI weights), pretrained=True | visual.transformer.resblocks.5, .11, .17, .23; visual | 1,024, 1,024, 1,024, 1,024, 768 | thingsvision model-specific transforms | Blocks: mean of patch tokens excluding CLS; visual: joint image–text embedding (ln_post + projection) |
| CLIP-RN50 | Vision–language model with modified ResNet-50 image encoder (Radford et al., 2021) | thingsvision custom model clip, variant RN50 (OpenAI weights), pretrained=True | visual.layer1, visual.layer2, visual.layer3, visual.layer4, visual | 256, 512, 1,024, 2,048, 1,024 | thingsvision model-specific transforms | Stages: global average pooling; visual: attention-pooled, projected embedding |
| ResMem | ResNet-50-based network trained to predict image memorability (Needell & Bainbridge, 2022) | resmem package, ResMem(pretrained=True) | features.layer1, .layer2, .layer3, .layer4; features | 256, 512, 1,024, 2,048, 1,000 | Resize 256 × 256, centre crop 227, ImageNet mean/s.d. normalization | Layers: global average pooling; features: flattened output |
